# Development and application of a cultivation platform for mammalian suspension cell lines with single-cell resolution (MaSC)

**DOI:** 10.1101/2020.07.14.202036

**Authors:** Julian Schmitz, Sarah Täuber, Christoph Westerwalbesloh, Eric von Lieres, Thomas Noll, Alexander Grünberger

## Abstract

In bioproduction processes cellular heterogeneity can cause unpredictable process outcomes or even provoke process failure. Still, cellular heterogeneity is not examined systematically in bioprocess research and development. One reason for this shortcoming are the applied average bulk analyses, which are not able to detect cell-to-cell differences. In this work we present a microfluidic tool for single-cell cultivation of mammalian suspension cells (MaSC). The design of our platform allows long-term cultivation at highly controllable environments. As model system CHO K1 cells were cultivated over 150 h. Growth behavior was analyzed on single-cell level and resulted in growth rates between 0.85 – 1.16 d^-1^, which are comparable to classical cultivation approaches such as shake flask and labscale bioreactor. At the same time, heterogeneous growth and division behavior, e.g., unequal division time, as well as rare cellular events like polynucleation or reversed mitosis were observed, which would have remained undetected in a standard population analysis based on average measurements. Therefore, MaSC will open the door for systematic single-cell analysis of mammalian suspension cells. Possible fields of application represent basic research topics like cell-to-cell heterogeneity studies, clonal stability, pharmaceutical drug screening and stem cell research, as well as bioprocess related topics such as media development and novel scale-down approaches.

## Introduction

Traditionally, analytical investigations in bioprocess research and development have been realized by bulk measurements. Consequently, every discovery and technical advancement is based on averaged population values determined over millions of cellular events (Templer and Ces 2008; Wang and Bodovitz 2010). As a result, individual behavior of non-conform acting cells in a genetically identical population remains masked behind average values which therefore do not depict every cell’s nature (Lecault et al. 2012; Lindström and Andersson-Svahn 2010; Yin and Marshall 2012). In the last years these cellular heterogeneities have received increasing awareness in consideration of being the origin of manifold inconsistencies in bioprocess related affairs such as viability and productivity (Delvigne and Goffin 2014). Moreover, in matters of biopharmaceutical production, heterogeneous behavior concerning overall growth or product yield can have severe impact on the process robustness and reproducibility. Of particular interest are properties like loss of productivity (Le et al. 2012), single-cell growth behavior and the occurrence of dormant cells during the bioprocess (Grünberger et al. 2015), as well as apoptotic and necrotic processes inside the bioreactor (Grilo and Mantalaris 2019). Certainly, conventional process analytical technology can detect the outcome of these phenomena, but has no opportunity for analyzing and understanding them, due to its lack of single-cell resolution and defined environmental conditions.

As a consequence of the rising interest in the behavior of individual cells during bioproduction applications, the number of single-cell analysis tools has increased over the last years (Schmitz et al. 2019). First approaches based on flow cytometry allow insights into single-cell behavior and population heterogeneity. However, dynamic investigation of single cells is not feasible. Therefore, especially approaches utilizing PDMS-based microfluidic devices became popular over the last years because of the steadily evolving microfabrication and ongoing progress in the field of microfluidics (Halldorsson et al. 2015). Their potential to trap cells in diverse featured microfluidic structures in combination with live cell imaging results in a high spatial and temporal resolution of single-cell behavior (Grünberger et al. 2014; Rowat et al. 2009). Depending on the systems design, microfluidic approaches provide high throughput analysis with hundreds of replicates on one microfluidic device. In carefully designed setups, a high level of environmental control can be achieved and even dynamic changes in cultivation conditions, reagent concentration, or nutrient supply can be realized (Dettinger et al. 2018; Marques and Szita 2017; Täuber et al. 2020).

Microfluidic single-cell analysis and cultivation have successfully been applied to examine bioprocess relevant questions of industrially relevant microbial host systems such as the influence of heterogeneity on cellular growth and production, the influence of specific media components, or bacterial co-culture approaches (Binder et al. 2016; Burmeister et al. 2018; Unthan et al. 2014). Lately, microfluidic single-cell applications likewise made their entry in the field of cell culture (Mehling and Tay 2014). Examples of applications are transfection studies (Raimes et al. 2017) and cell line development (Bjork et al. 2015), cell interaction studies (Li et al. 2016), drug screening (Du et al. 2016), and stem cell research (Luni et al. 2016) where single-cell analysis expanded the previous state of knowledge. In general, microfluidic cultivation devices can be discriminated by their way of capturing and retaining single cells into traps, wells, or chambers (Fig. 1). These setups have different system specifications that enable different applications. Microfluidic traps (Fig. 1 A) have been used for e.g., viability assays and morphology studies focusing on single-cell behavior and not on microcolony growth (Di Carlo et al. 2006; Wheeler et al. 2003). The cells are located inside the medium stream which allows convective supply of nutrients during the experiment but also holds the risk of shear stress. Since cells are actively retained inside the trap, the cultivation of adherent as well as cells growing in suspension is realizable. In comparison to traps, the supply of nutrients in cultivation wells (Fig. 1 B) is diffusive and convective as cells are located inside the wells and are perfused across. Therefore, cultivation of suspension cells is not feasible without washing them out frequently. Depending on well diameter microcolony growth can be observed which makes wells especially applicable for cell spreading, proliferation, differentiation, and cell interaction studies (Karakas et al. 2017; Lin et al. 2015). Cultivation chambers combine the advantages of traps and wells for the cultivation of cells (Fig. 1 C). Flow velocities inside the chambers are very low, meaning a large part of the nutrient supply takes place by diffusion and the cells are not exposed to significant shear stress. Although cultivation of adherent as well as suspension cells is conceivable, so far only adherently growing cells have been cultivated in chambers. Due to their characteristics which make long-term growth feasible, chambers were applied in growth studies and drug screening (Gao et al. 2013; Kolnik et al. 2012).

**Fig. 1:**
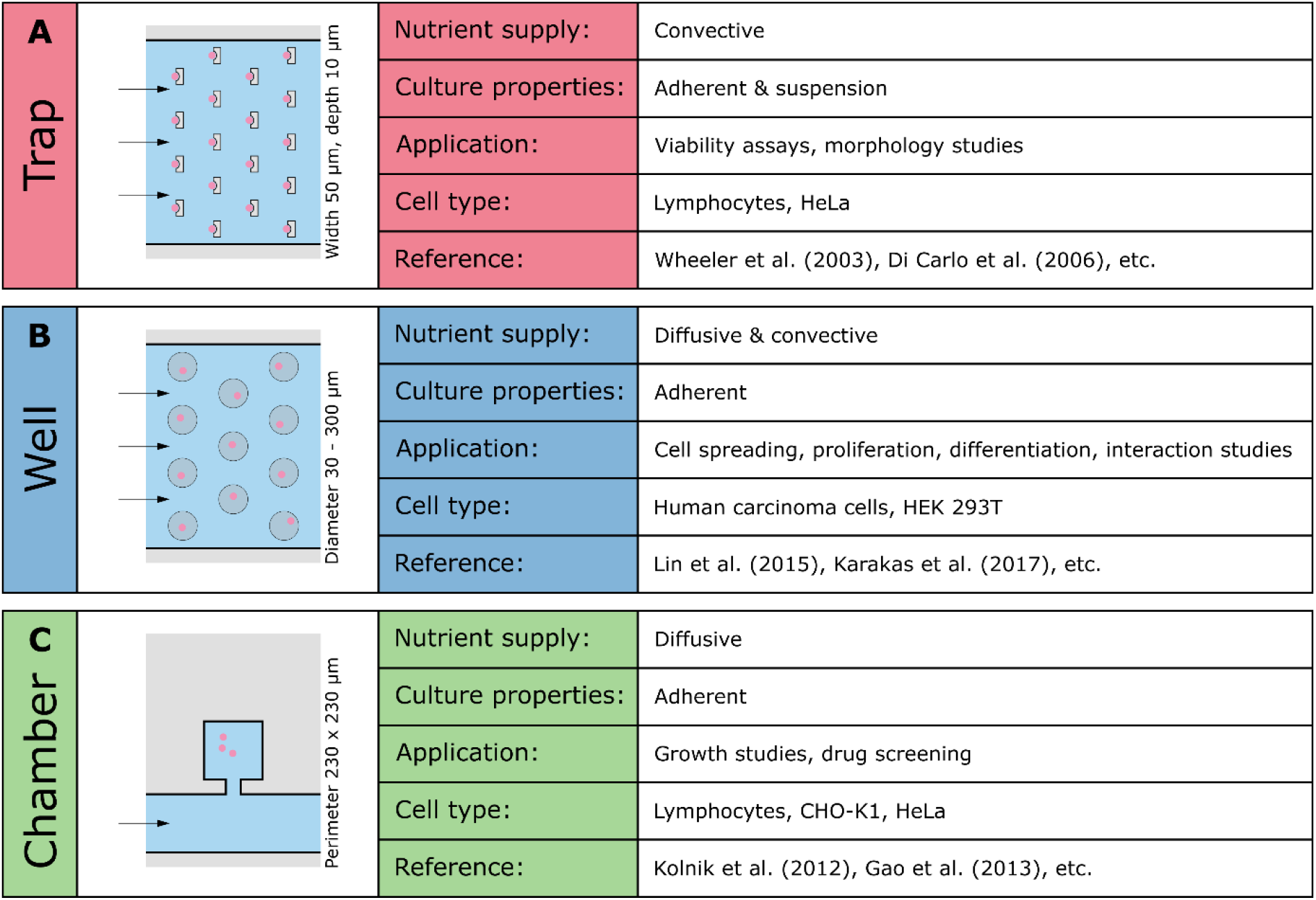
Examples of microfluidic cultivation devices in the field of mammalian cell culture categorized into the underlying cell retention principle. Traps, wells, and chambers differ in the way cells are retained and supplied with nutrients, offer appropriate cultivation surfaces for either adherent or suspension cell lines and support different applications. For every device examples of utilized cell types as well as representative studies are listed.

Until now microfluidic single-cell applications for mammalian cell lines are limited to analytical questions in basic and applied research but do not systematically address single-cell growth, media development, and process engineering in terms of bioprocess research and development (Schmitz et al. 2019). In contrast to the already existing microbial applications, almost all mammalian approaches to study individual cellular behavior do not offer the opportunity to examine growth characteristics on single-cell level for appropriate cultivation times. This is due to their lack of sufficient cell trapping, insufficient nutrient supply, and lack of live cell imaging. As a proof-of-concept, first studies investigating long-term single-cell growth of adherent CHO and HeLa cell lines have been reported and showed comparable growth between microfluidic and lab-scale cultivation (Kolnik et al. 2012). Nevertheless, these systems were not applied for systematic growth studies of mammalian cell lines with full spatio-temporal resolution in the context of dynamic single-cell studies. An even larger drawback is the exclusive application of adherent growing cells. Since most bioproduction processes rely on in suspension growing cells (Wurm 2004), results from single-cell cultivation of adherent growing cells are of limited relevance for this application. Consequently, single-cell cultivation of mammalian cells is not only hampered by the unavailability of appropriate setups, but also by non-existent studies of suspension cell lines in microfluidic applications.

In this work we present a PDMS-glass-based microfluidic setup for long-term single-cell cultivation of CHO cells growing in suspension. This setup allows simultaneous analysis of growth characteristics and cellular heterogeneity under highly controlled environmental conditions. Using soft lithography microfluidic chips were fabricated, which offer the possibility to capture and track single mammalian cells in isolated cultivation chambers. Adjacent perfused supply channels enable constant delivery of nutrients and removal of metabolic by-products, due to diffusive processes without exposing the cells to shear stress. Sufficient mass transfer was verified by computational fluid dynamics (CFD) simulations. Growth of CHO K1 cells was investigated and evaluated on single-cell level. The results show reproducible growth behavior, which is comparable to cultivations in standard bioreactors and shake flasks, making MaSC an attractive single-cell cultivation device.

## Material and Methods

### Microfluidic device fabrication

Using two-layer photolithographic techniques described previously (Grünberger et al. 2013a), a silicon wafer was produced as mold for PDMS (polydimethylsiloxane) chip fabrication. PDMS base and curing agent (Sylgard 184 Silicone Elastomer, Dow Corning Corporation, USA) were mixed in a ratio of 10:1, poured onto the silicon wafer, degassed by means of a desiccator, and cured for 2 hours at 80 °C. Afterwards the chips were cut, inlets and outlets were punched with a biopsy punch (Reusable Biopsy Punch, 0.75mm, WPI, USA), and particles were removed from PDMS chip and glass substrate (76 x 26 x 1 mm microscope slides, VWR International GmbH, Germany) via isopropanol washing and drying with pressurized air. To covalently link chip and glass, surfaces were activated by oxygen plasma (Femto Plasma Cleaner, Diener Electronics, Germany) and bonded to each other. Subsequently, the microfluidic device was incubated at 80 °C for 1 min to enhance bonding.

### Computational Fluid Dynamics

Detailed CFD simulations were performed to model flow profiles within cultivation chambers and supply channels as well as to analyze nutrient supply inside the microfluidic device. The computational model used in this study has been described in detail previously (Westerwalbesloh et al. 2015; Westerwalbesloh et al. 2017). The model geometry contains only one chamber and parts of the adjacent supply channels, because the conditions within each chamber are assumed to be independent of its position on the microfluidic device. All CFD simulations were performed using COMSOL Multiphysics Version 5.4 (COMSOL AB, Sweden). The flow field was determined by solving the steady-state Navier-Stokes equations for creeping (Stokes) flow of an isothermal, incompressible and Newtonian liquid with the properties of water (density 993.22 kg m^-3^, viscosity 6.96 × 10^-4^ Pa s (interpolated for 37°C (Comesaña et al. 2003)). The no-slip condition was used at the PDMS and glass walls. Laminar inflow with a rate of 8.33 × 10^-12^ m^3^ s^-1^ was specified at each of the supply channel inlets, while the outlet condition was a reference pressure of 0 Pa. The glucose concentration field was calculated by solving the general steady state diffusion-advection equation with a diffusion coefficient of 8.5 × 10^-10^ m^2^ s^-1^. Adsorption at the glass and PDMS walls was neglected. Due to the concentration of glucose being below 45 mol m^-3^ it could be assumed that the flow field is not influenced by the glucose concentration, and therefore flow and mass transfer were solved sequentially. In the model the chamber is populated with cells (171 cells in a filled chamber on a typical microscope image). Every cell is assumed to take up glucose with a rate of 3800 nmol per 10^6^ cells and day (Nolan and Lee 2011), which leads to a volumetric uptake rate of 2.4 × 10^-2^ mol s^-1^ m^-3^, distributing the cellular uptake homogeneously over the growth chamber. The geometry of the cells themselves is not explicitly included as shown before (Westerwalbesloh et al. 2017). The glucose concentration provided in the growth medium is assumed to be high enough to yield a constant (maximal) uptake rate in the model, independent of the exact local glucose concentration. The computational geometry was discretized using 106,176 rectangular elements. A finer mesh with 847,000 elements yielded the same results, indicating mesh independence of the solution (data not shown). Quadratic functions were used to calculate velocity profiles and concentrations and linear functions for the pressure.

### Cell culture and medium

Cultivation of Chinese hamster ovary (CHO) K1 cells was performed in commercially available cell culture medium (TCX6D, Xell, Germany), supplemented with 8 mM glutamine. Cultivation conditions for pre-culture were 37 °C, 5 % CO_2_, 80 % humidity, and 180 rpm in TubeSpin^®^ Bioreactor 50 (TPP^®^, Switzerland) with a cultivation volume of 15 mL. CHO K1 cells were subcultured every 3^rd^ respectively 4^th^ day and viable cell density was adjusted to 1.5 x 10^5^ cells mL^-1^. For microfluidic single-cell cultivation, fresh medium was mixed with conditioned medium in a ratio of 1:1. For the generation of conditioned medium, cells were cultivated according to the upper protocol, the required volume of cell suspension was centrifuged at first and the resulting supernatant was additionally sterile filtrated using a 0.2 μm cut-off filter (Filtropur S 0.2, Sarstedt AG & Co KG, Germany).

### Microfluidic single-cell cultivation

Single-cell cultivation in microfluidic devices was performed on an automated inverted microscope (Nikon Eclipse Ti2, Nikon Instruments, Germany), enabling high-throughput time-lapse microscopy. Cultivation temperature was kept constant at 37 °C by using a microscope incubator system (Cage incubator, OKO Touch, Okolab S.R.L., Italy). An additional CO_2_ incubation chamber (H201-K-FRAME GS35-M, Okolab S.R.L., Italy) enabled constant CO_2_ atmosphere of 5 % (95 % compressed air). Monitoring growth and morphology was achieved by live cell imaging: Time-lapse images of relevant positions were taken every 20 min (NIS Elements AR 5.20.01 Software, Nikon Instruments, Germany) applying 40x objective with phase contrast microscopy.

Cells were seeded into cultivation chambers by manually flushing the microfluidic chip with cell suspension until loading of the chambers was sufficient. Medium supply was realized by low pressure syringe pumps (neMESYS, CETONI, Germany) and single-use syringes (Injekt^®^ 10 ml, B. Braun Melsungen AG, Germany) connected to the chip via PTFE tubing. During cultivation, medium was constantly perfused through the cultivation device with a flow rate of 2 μL min^-1^.

### Image evaluation and growth analysis

Cell number was manually counted on time-lapse images every 10 h to evaluate growth behavior. The colony growth rate μ was estimated graphically by determining the slope of the linear regression from the resulting semi-logarithmical plot using OriginPro (OriginPro 2020 9.7.0.188, OriginLab Corporation, USA). Under the assumption of exponential colony growth, equation (1) was applied to convert μ to doubling times t_D_

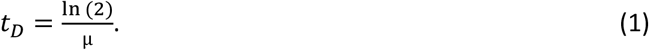

Since exponential growth of mammalian cell lines can be described by a geometric sequence, respective mean values for t_D_ and μ were determined using the geometric mean (Phoenix 1997). In comparison to the arithmetic mean, which is especially suitable for linear sequences, the geometric mean is less prone to the influence of outliers in a broad distribution of values. For quantification of single-cell doubling time distribution, single cells were tracked manually and the time span between two cell divisions was determined. Single-cell area was analyzed using ImageJ 1.52p (Schindelin et al. 2012). For determination of area growth, cellular contours on scaled microscope pictures were manually retraced. Cellular area was subsequently determined and plotted against cultivation duration. Analog to colony growth rates, these charts were used to determine area related single-cell growth rates.

## Results and discussion

### Device design and working principle

Derived from a previously introduced microbial system, the microfluidic device presented in this work was optimized for single-cell cultivation of mammalian cell lines in high throughput, with high spatio-temporal resolution, and at highly controlled environmental conditions. One microfluidic device consists of four independent cultivation arrays with one inlet and one outlet each for environmental interfacing (Fig. 2 A). Incorporating two times 30 cultivation chambers with dimensions of 200 x 200 μm, each array is composed of four parallel running supply channels with a width of 200 μm (Fig. 2 B). Two channels are connected to one intermediate chamber. 50 x 50 μm entrances link the cultivation area with the supply unit of the array. Growth chambers and entrances are approx. 8 μm high, while supply channels are designed with a height of 16 μm. The difference in height of the structures is intended to restrict flow from the channel into the chambers and facilitate cell retention inside the chamber. Likewise, the narrow chamber entrances improve withholding seeded cells. As a result of the limited height, cellular growth inside the chamber is restricted to monolayer growth. Resulting from its dimensions, the array’s total volume sums up to approx. 2.3 x 10^8^ μm^3^, equaling 230 nL. Due to the chip’s fabrication using soft-lithography, silicon wafers with varying channel and chamber heights can easily be fabricated to adapt chip dimensions to specific scientific question and cell lines.

**Fig. 2:**
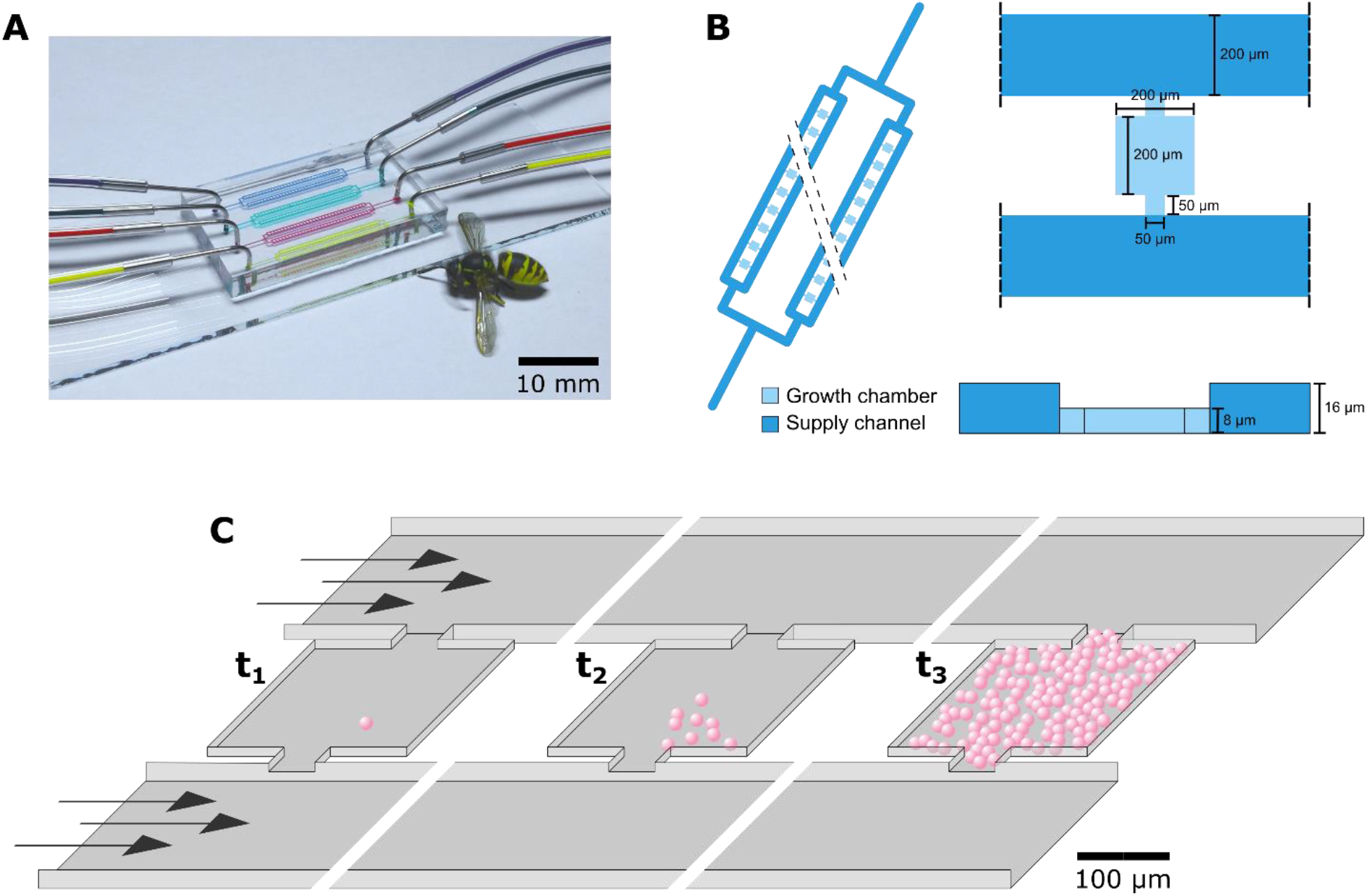
Design and working principle of MaSC. (A) Microfluidic PDMS-glass cultivation device for single-cell cultivation of mammalian suspension cell lines. (B) Schematic figure of the cultivation device: Each cultivation array holds two times 30 cultivation chambers with an area of 200 x 200 μm and a height of approx. 8 μm. The adjacent supply channels are twice as high and show a width of 200 μm. (C) Three-dimensional illustration of the microfluidic device at different stages of the cultivation. t_1_: A single cell is seeded into the cultivation chamber. t_2_: The starter cell begins to proliferate and exponential growth starts. t_3_: Approaching the end of a cultivation the chamber is filled with up to 170 cells.

To capture single cells inside the cultivation chambers, cell suspension is flushed through the supply channels and cells randomly enter the chambers. Since cell seeding is not actively controllable, loading efficiency meaning the percentage of seeded and empty chambers is directly connected to cell density of the inoculum. Here, concentrations around 30 x 10^5^ cells mL^-1^ proved to be a reasonable compromise between loading efficiency and cell number per loaded chamber. If cell density is lower, only a few chambers are loaded with cells. If cell density is too high, the starting cell number exceeds one to five cells per chamber promoting non-isogenic microcolonies.

Once enough cultivation chambers are loaded with cells, constant perfusion of the array is initiated through the chip’s inlet by syringe pumps (Fig. 2 C). With a steady flow of 2 μL min^-1^ cells are constantly provided with fresh nutrients while metabolic by-products are washed out through the outlet. This flow rate provides an exchange of the entire microfluidic device’s volume about 10-times every minute, which assures constant environmental conditions over the whole cultivation course. Starting the cultivation with a small number of cells, the final cell number per cultivation chamber can reach up to 170 cells. Further exponential proliferation is only limited by the chamber’s restricted volume of 3.2 x 10^-7^ mL.

### Device characteristics and functionality

CFD simulations were performed for detailed evaluation of fluid profiles inside the supply channels and cultivation chambers as well as the nutrient related cultivation conditions. With the presumed flow rate of 0.5 μL min^-1^ for each supply channel (4 channels per inlet), a parabolic velocity profile and a maximal velocity of 30 x 10^-4^ m s^-1^ is observed inside the channels (Fig. 3 A). Due to the symmetric layout of the design as well as the narrow entrance, fluid flow inside the cultivation chamber is too slow to cause perturbations (Fig. 3 B). Thus, exchange of nutrients and by-products is predominantly diffusive. To examine whether cells inside a completely filled cultivation chamber can possibly experience limiting glucose concentration, another simulation was performed. With 171 cells per chamber and the specific glucose uptake rate of CHO K1 cells, glucose concentration inside the chamber does not drop below 43.8 mmol L^-1^ in a steady-state (Fig. 3 C). Furthermore, a medium switch was simulated to determine the duration until full equilibrium from zero glucose to a cultivation sufficient glucose concentration of approx. 45 mmol L^-1^ is accomplished. The simulation indicates that after 25 s the cultivation chamber already shows half-maximal glucose concentration, whereas 99,9 % of the medium’s glucose concentration is reached after approx. 150 s homogeneously throughout the whole chamber (Fig. 3 D).

**Fig. 3:**
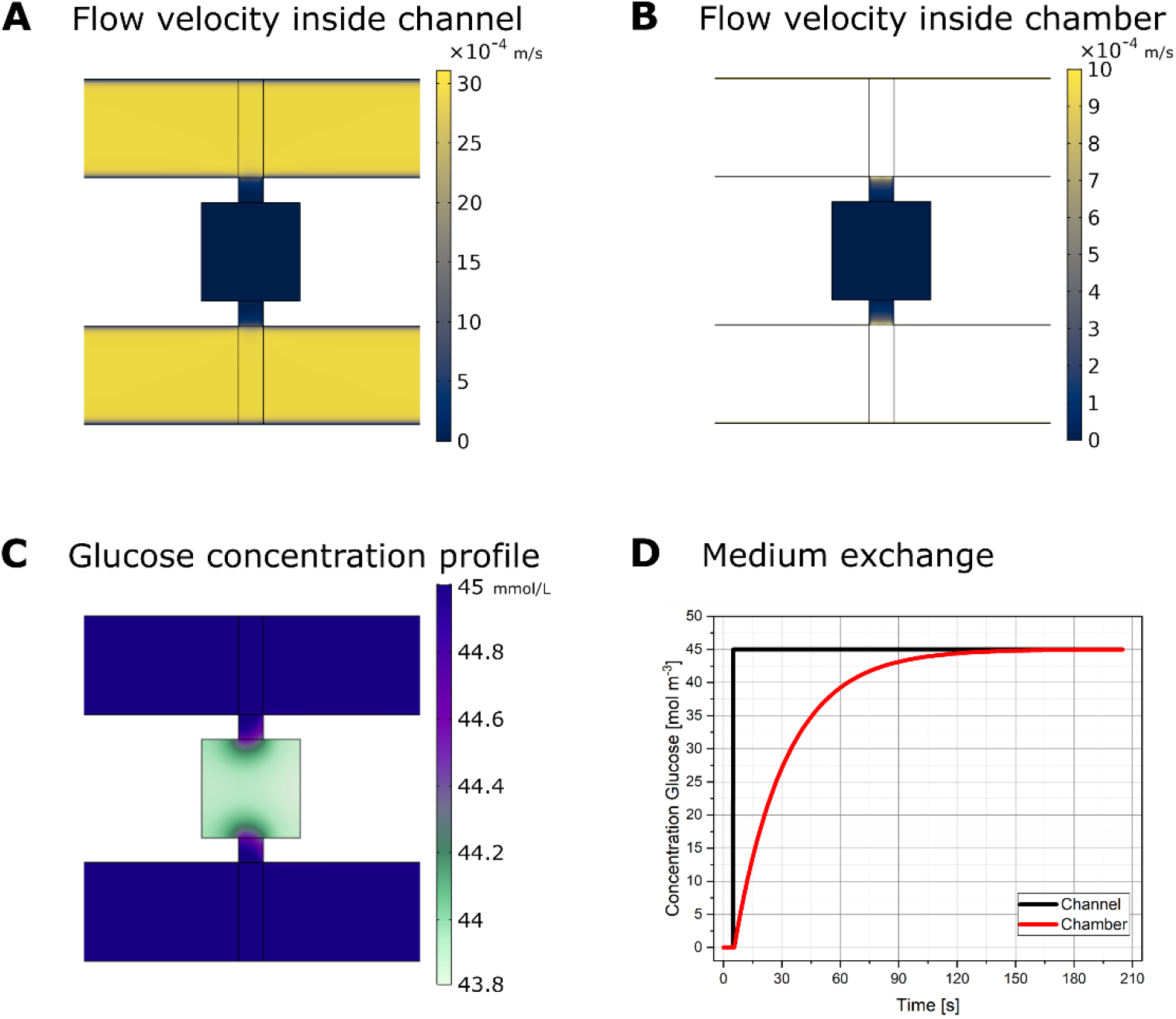
CFD simulations of flow and nutrient profile inside microfluidic device. (A) Flow velocity inside supply channels. (B) Flow velocity inside representative cultivation chamber. (C) Glucose concentration profile assuming a steady-state at a cell number of 171 cells inside the chamber. (D) Glucose concentration inside the supply channels and cultivation chamber after quick medium change from non-glucose to 45 mmol L^-1^ glucose.

### Mammalian single-cell cultivation

Since CHO cell lines are the most frequently used cells in biopharmaceutical bioproduction processes (Walsh 2018), CHO K1 cells were used as a model system for the developed microfluidic device. For process near growth conditions, a commercially available cell culture medium, optimized for growth and production with CHO cell lines, was employed.

In three consecutive experiments precultured cells were seeded into the microfluidic device. Since cell loading is a mere statistical process, the number of loaded cultivation chambers varied in a range of 15 to 25 chambers. The number of cells in each chamber after loading varied between one and seven cells. As the medium exchange rate in the chambers is very high and to mimic the substrate and metabolite situation in standard bioreactors, a 1:1 mixture of fresh and conditioned medium was used as cultivation medium.

Cell growth in the chambers was analyzed on colony level as well as on single-cell level. Here, colony growth rate μ_colony_ refers to the characteristic average values obtained from the cell number of each colony. In contrast, single-cell doubling times t_D, single-cell_ refer to characteristic values obtained from single cells. In addition, single-cell area growth rates μ_area_ are obtained from single-cell area development. Under the assumption of an exponential growth behavior, each growth rate μ can be converted into the respective doubling time t_D_ and vice versa using equation (1).

### Colony growth analysis

During the experiment cells grew exponentially until the whole chamber was filled. The time-lapse sequence of Fig. 4 A illustrates an example of a growing microcolony. Random migration of cells inside the cultivation chambers could be observed (Video S1). After six to seven days a maximal cell number of approx. 150 per chamber is reached. At this state, cellular boundaries blur, since dividing cells are squeezed so tightly that individual cell membranes cannot be identified microscopically anymore. A cell number of 150 represents a cell density of 4.8 x 10^8^ cells mL^-1^ inside the cultivation chamber which is similar to the cellular density of human tissue (Skylar-Scott et al. 2019). Protuberances of cellular membrane is observed during cultivation and caused by microvilli, extracellular matrix, endocytosis, and vesicle formation. Right before cell division, cells become sphere-shaped and smooth until cytokinesis took place and the originated daughter cells start to show protuberances again (Fig. 5 A).

**Fig. 4:**
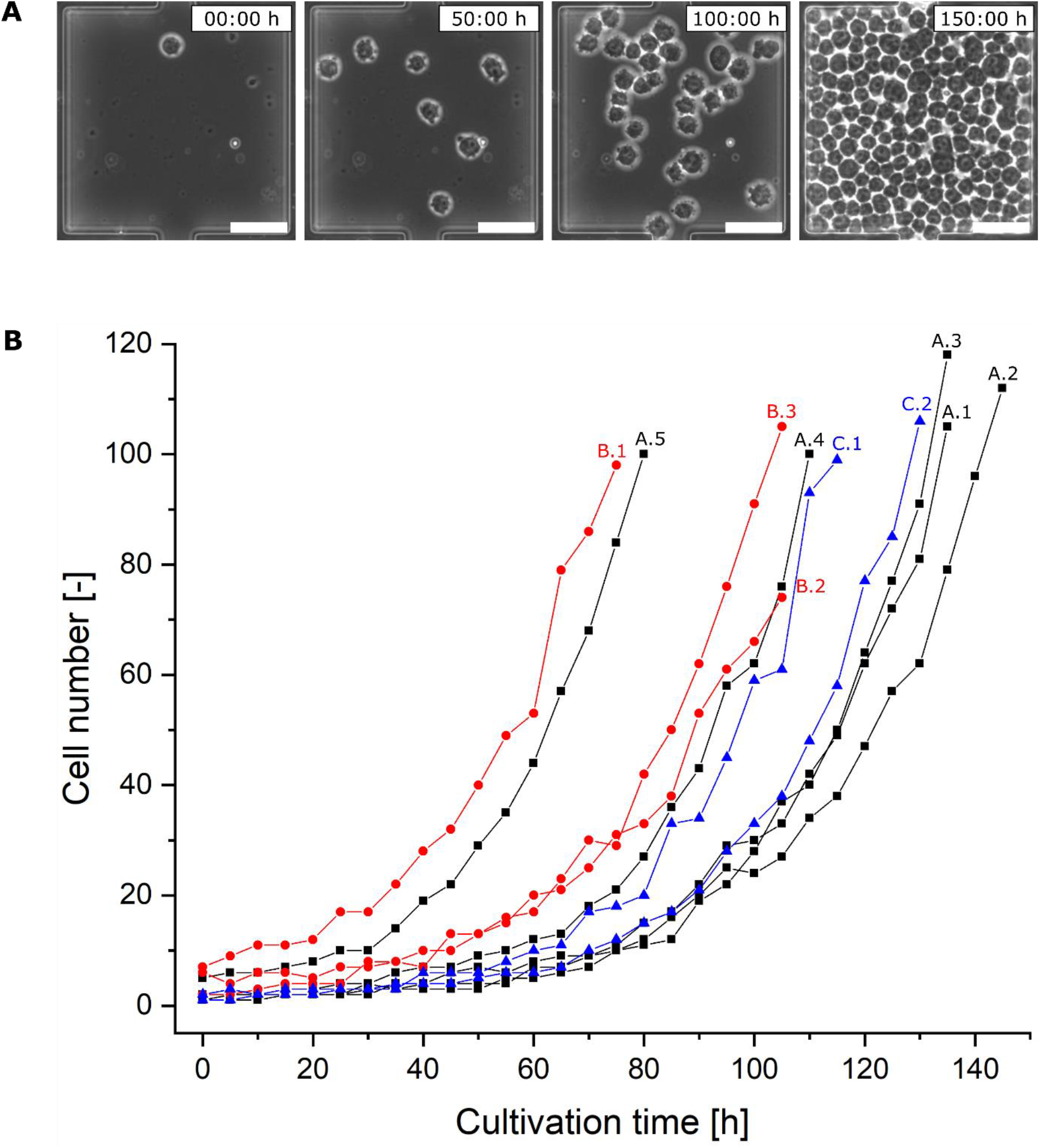
Colony growth of CHO K1 cells at constant environmental conditions. (A) Time-lapse image sequence showing microfluidic growth of one initial CHO K1 cell to a microcolony over an experimental duration of 150 h (scale bar 50 μm). (B) Colony growth of CHO K1 microcolonies. Here, ten selected chambers from three technical replicates are shown. Black squares (A.1 – A.5), red dots (B.1 – B.3), and blue triangles (C.1 – C.2) represent one technical replicate each. Depending on starting cell number and cell cycle distribution, exponential growth starts slightly shifted between the microcolonies.

**Fig. 5:**
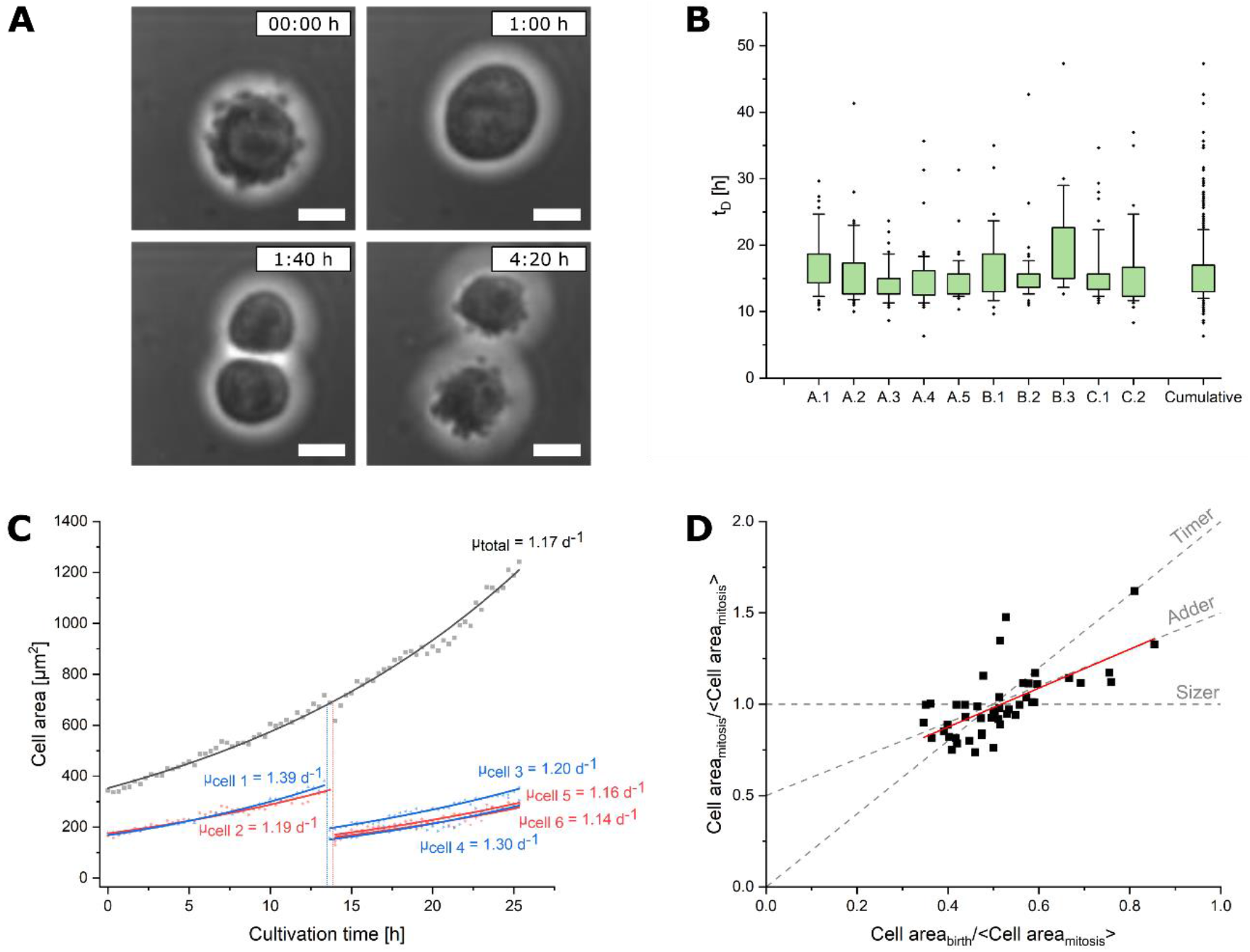
(A) Cell division event for illustration of the accompanying cellular morphological changes (scale bar 10 μm). (B) Distribution of single-cell doubling times of the microcolonies depicted in Fig. 4 complemented with a cumulative t_D_ distribution over all doubling events. The colored segment marks the interquartile range from 25% to 75 %. The whiskers represent the 10 % and 90 % percentile and the tilted squares mark rare cellular events. (C) Determination of area growth rates of single cells from colony A.5. The analysis started with the origin of cell 1 and cell 2 from one mutual progenitor cell and was conducted for 25 h in intervals of 20 min. Depicted are the colony’s total cell area and the individual cells’ areas. By exponential fit specific growth rates were determined from increasing cell area. (D) Cell size homeostasis analysis of single cells from colony A.1. To which of the conventional cell size homeostasis models (timer, adder, and sizer) applies to CHO K1 cells, the correlation between cell area at birth and cell area at mitosis was analyzed. For clearness, all area data was normalized by the average cell area at mitosis. The trend lines (dashed grey) show the expected trends in case of timer, adder, and sizer behavior.

Depending on seeding cell density within the chambers, colony growth curves rise slightly shifted in the beginning, but initial cell number does not have a notable influence on growth progression of each microcolony (Fig. 4 B). Most likely, cell cycle distribution influences time until first cell division starts, since cells were randomly distributed across cell cycle phases before loading the microfluidic device. This can result in a delay of up to 20 h for cells being in G1 phase compared to cells in G2 phase, which start mitosis very rapidly. A lag-phase for chambers with low initial cell number seems unlikely, given that semi-logarithmic growth curves show exponential growth right from the experimental start (Fig. S1). Linear fits show similar colony growth rates between 0.75 d^-1^ and 0.95 d^-1^ for the colonies presented here (Tab. S1).

### Single-cell growth analysis

Besides evaluating microcolony growth, MaSC explicitly enables single-cell data acquisition (Fig. 5 A). Therefore, we determined single-cell doubling times and single-cell area development to analyze cellular division behavior throughout the microcolony. As can be seen in Fig. 5 B, the distribution of single-cell doubling times t_D, single-cell_ underlies certain variabilities, especially the distribution width differs and rare cellular events can be identified; most of them show an extended doubling time longer than 25 h, only few fall below 10 h. Looking at the respective mean doubling time they alternate between 14.3 and 19.6 h whereas the cumulative determined mean doubling time is 15.9 h (Fig. S2). Looking at colony A.5, B.1, and B.3 (Fig. 5 B) shows that starting cell number influences t_D_ only marginal since each colony started with a cell number ≥5 and broadness as well as mean t_D, single-cell_ differ clearly. Since correlation coefficient for starting cell number and mean t_D_ is R = 0.394, only a weak trend can be assumed (Fig. S3).

Fig. 5 C shows the single-cell area development of two cells originated from a mutual progenitor cell. The area increase shows small variations, the corresponding μ_area_ differ between 1.14 and 1.39 d^-1^. Noteworthy, second generation cells originated from the slower initial cell both show slightly lower area growth rates than those originated from the faster initial cell. Comparing the area growth rate with the corresponding doubling times of the analyzed cells shows no correlation between μ_area_ and t_D, single-cell_ (Fig. S4). One explanation might be the continuous secretion of vesicles, which makes the cellular area a continuously fluctuating property and thus the exact determination of the cell area difficult. Another more relevant reason lies in the dependency of cellular growth on cell cycle phases combined with the cell’s mechanism to ensure constant cellular size over generations. In general, there are three different models that describe how cells can reach size homeostasis. The sizer-mechanism controls the beginning of mitosis by a size threshold, for the adder-mechanism the cell size increases by a constant amount independent of initial cell size, and with the timer-mechanism cells grow exponentially for a constant duration (Cadart et al. 2018; Vuaridel-Thurre et al. 2020). To determine if this correlation applies on the area development of CHO K1 cells, we compared the cell area of single cells at their birth and at their mitosis. For better clearness, the data in Fig. 5 D were normalized by the average cell area at mitosis. Since sizer-cells undergo mitosis at a consistent cell size independently from their size immediately after mitosis, this mechanism can be illustrated by a horizontal line. A line with a slope of 1 and an intercept of 0.5 represents cells with adder-behavior while timer-cells be in consistent with a line through origin and a slope of 2 (assuming exponential cell growth). Comparing the single-cell data with the expected trends an adder-mechanism to assure size homeostasis can be identified for the here analyzed CHO K1 cells. Additionally, a correlation between cell area_birth_ and the ratio of cell area_mitosis_ and cell area_birth_ casts a timer mechanism in doubt (Fig. S5). This behavior is in consistency with already analyzed mammalian cells, which were analyzed in an microfluidic flow-through system (Cadart et al. 2018).

Growth rates obtained from single-cell level are noteworthy higher than those obtained from colony level (Tab. S1). Here, the main reason for this discrepancy are technical insufficiencies of the developed microfluidic setup, as occasionally single cells can escape the cultivation chamber by random movement reducing the apparent colony growth rate. Such events can be identified as bends in some colony’s growth curves (Fig. 4 B). Therefore, the here determined colony growth rates underestimate growth rates which would be obtained without the described cell loss.

Consistencies between microfluidic cultivation and larger cultivation scales have already been described before by Kolnik and colleagues, who compared growth of adherent CHO cells in a microfluidic device with 6-well-plate cultivation (Kolnik et al. 2012). With MaSC the obtained growth rates of CHO suspension cells were similar to classic shake flask or lab-scale bioreactor cultivations (Schmitz et al. in preparation). Transferring common cultivation scales into microfluidic devices does not always show comparable growth behavior, since steady perfusion leads to constant wash-out of secreted molecules and beneficial factors (Taheri-Araghi et al. 2015). Preliminary tests showed that CHO K1 cells do not grow in fresh medium (data not shown). For this reason, we supplemented commercial medium with already conditioned one in our study. Other examples like the cultivation of the microbial production hosts such as *C. glutamicum* showed significantly higher growth rates in microfluidic single-cell cultivation compared to lab-scale cultivations have been achieved (Fritzsch et al. 2013; Grünberger et al. 2013b), which most likely result from the growth-promoting environmental conditions concerning nutrient supply inside the applied microfluidic devices.

### Cell-to-cell heterogeneity and rare cellular events

During CHO K1 single-cell cultivation described above, various examples of heterogeneous behavior between isogenic cells have been observed. Considering the rare cellular events of Fig. 5 B, variations in cell division happen in every cultivation but only single-cell analysis makes them visible. In the following some of these outliers with t_D_ > 25 h were analyzed more closely on single-cell level.

Originating from a common mother cell (G0), the subsequent doubling time of the daughter cells (first-generation G1) varied strongly (Fig. 6 A & Video S2). While one daughter cell (depicted in red) grew in cell volume for another 18 h until it finally underwent cell division, the other daughter cell (depicted in green) and its progenies (second-generation G2) finished two rounds of cell division.

**Fig. 6:**
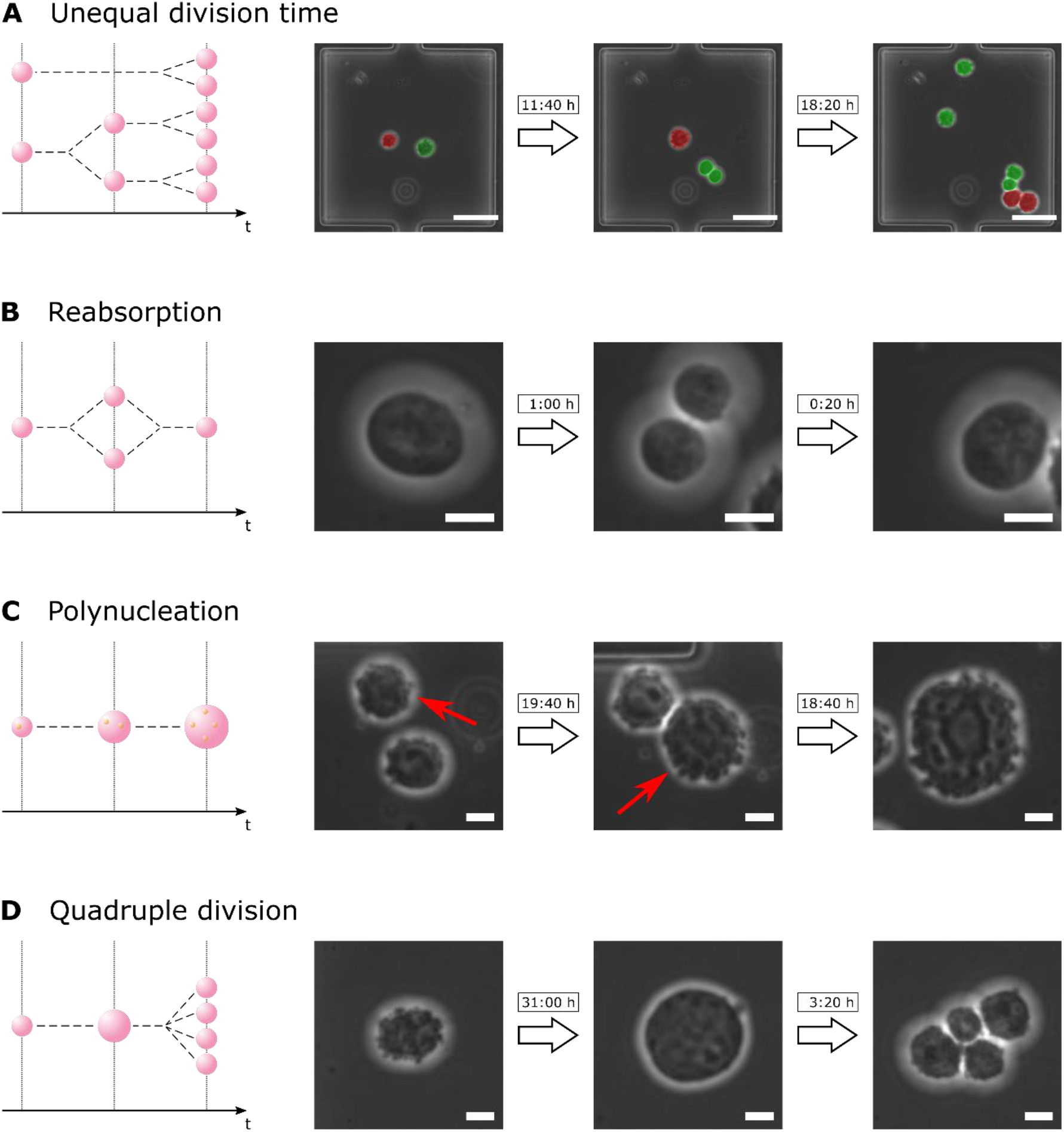
Examples of cell-to-cell heterogeneity and rare cellular events during mammalian single-cell cultivation. (A) Genetically identical cells differ in their doubling time by 18 h (scale bar 50 μm). (B) One G1 cell reabsorbs the second G1 cell immediately after cell division (scale bar 10 μm). (C) Absence of cell division leads to immense polynuclear cells (scale bar 10 μm). (D) One cell divides into four daughter cells which all are viable (scale bar 10 μm).

Delay in proliferation of single cells is not the only incident being observed. Cell division can also be aborted or even reversed in the final step of cytokinesis. After a performed division, one daughter cell reabsorbed the other (Fig. 6 B & Video S3). This observation in some respects contradicts the hypothesis of the point of no return in eukaryotic cell cycle. One explanation might be the occurrence of errors during mitosis, so that reabsorption acts like a safety mechanism against formation of defective progenies (Potapova et al. 2006). Few cells did not even start cytokinesis but nevertheless underwent DNA replication. Hence, huge polynuclear cells appear (Fig. 6 C & Video S4). Since they also occur in microcolonies where other cells show perfectly normal proliferation, detrimental environmental conditions are highly unlikely as cause for polynucleation. Surprisingly, in some cases these polynuclear cells eventually pass through cytokinesis. Sometimes, these cells divide into two daughter cells which randomly are polynuclear as well. In other cases, polynuclear cells divide into multiple mononuclear cells (Fig. 6 D & Video S5).

We assume that the – partly surprising – incidences being described in Fig. 6 are not caused by the microfluidic cultivation condition, but generally occur in CHO cell culture including bioreactors and biopharmaceutical processes and contribute to population heterogeneity.

### Future applications

The introduction of MaSC showed that long-term cultivation of mammalian suspension cells in chamber-based microfluidic devices is feasible. Besides growth analysis on colony level, also singlecell behavior can be examined and heterogeneity among individual cells can be revealed. Prospectively, MaSC may not only be applied for growth studies but also for systematical investigation of population heterogeneity, cellular transfection or cell-to-cell interactions in basic research (Fig. 7 A). In a more application-orientated way, our tool can be installed in different disciplines of (bio)pharmaceutical research (Fig. 7 B), ranging from first screening of potential drug components, the development of new cell lines for industrial production to stem cell research. With reference to bioproduction processes systematic design-of-experiment-based media development and process optimization represent possible fields of application for MaSC (Fig. 7 C). Eventually, single-cell cultivation of mammalian cells can even serve as starting point for bioprocess development in terms of scale-down approaches. However, beforehand comparability from microfluidic cultivation to shaker and lab scale bioreactor needs to be shown by in-depth validation studies to ensure transferability from single-cell level.

**Fig. 7:**
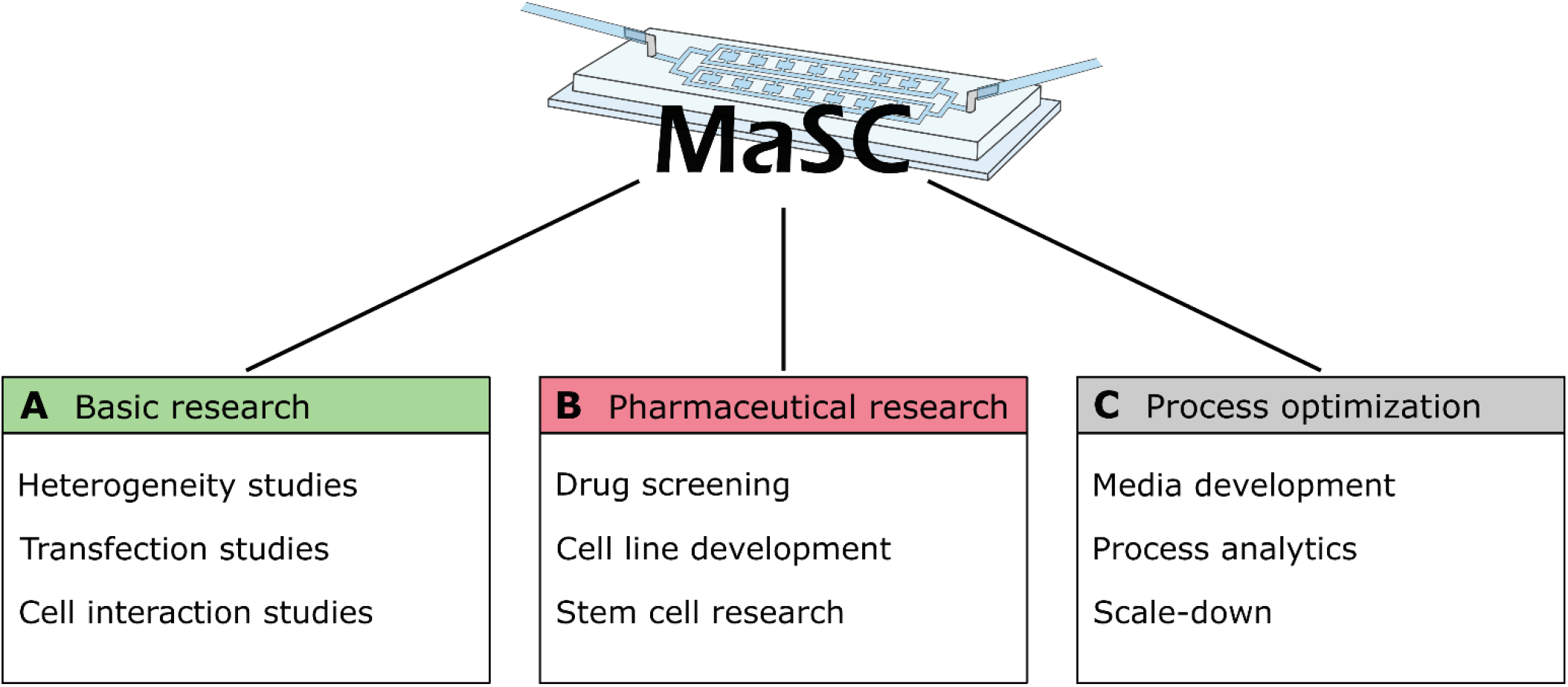
Examples of application fields for MaSC. (A) In basic research MaSC can be applied for heterogeneity studies, investigating transfection, and analyzing cellular interactions. (B) In pharmaceutical research especially drug screening, cell line development, and stem cell research approaches can be realized with MaSC technology. (C) In bioprocess research MaSC can simplify media development, function as process analytical tool, or prospectively be the starting point for scale-down approaches.

Before systematic application there are still technical aspects to improve and obstacles to overcome. First, better cell retention needs to be achieved by adapting the device’s design to prevent cell loss from the cultivation chamber due to movement of the cells. Likewise, cell loading efficiency must be increased. Concerning experimental data evaluation, automated image analysis appears worthwhile, as cell number, doubling times, and cell area were determined manually and therefore are error-prone and time-consuming.

## Conclusion

The MaSC platform presented in this study allows long-term cultivation of mammalian cells in suspension starting from a single cell and makes microfluidic systems available for industrially relevant biopharmaceutical applications. Microfluidic cultivation ensures controlled environmental conditions while live cell imaging enables spatio-temporal resolution of cellular behavior. Consequently, not only population dynamics but also individual single-cell events can be investigated. The device’s modular concept permits scalability from currently 60 individual cultivation chambers up to several hundreds to increase the amount of statistical data per run. Alongside with its other benefits, this aspect makes MaSC suitable for analytical studies of industrially relevant cell lines.

The focus of our study was establishing microfluidic single-cell cultivation for suspension cell lines. We achieved reproducible growth rates that are in accordance to shake flask or lab-scale bioreactor cultivations. Growth analysis was performed on microcolony as well as on single-cell level. Several, partly very surprising, incidences resulting in population heterogeneity have been observed on single-cell level and were investigated subsequently. This lays the foundation for systematic singlecell studies for suspension cell lines in the fields of basic research, cell line development and characterization, stability studies, and bioprocess development.

For the future it is interesting to extend the MaSC platform by adding tools that allow epigenetic and functional genomic analysis of individual cells and to include analysis of product quality when using recombinant production cell lines. This will enable an increased understanding of the full scope and consequences of the observed cellular heterogeneities and open the door for the development of stable (production) cell lines and homogeneous bioprocesses (Schmitz et al. 2019).

## Acknowledgements

Parts of this work were performed at the cleanroom facilities of the Department of Biophysics and Nanoscience as well as the Department for Physics of Supramolecular Systems and Surfaces at Bielefeld University. Cell culture maintenance was performed with help of the members of the Cell Culture Technology group at Bielefeld University. Preliminary single-cell cultivation experiments have been performed by Sebastian Perez Knoche and Lars Niklas Halle. The authors would like to thank all for help and support.

## Publication bibliography

Binder, Dennis; Probst, Christopher; Grünberger, Alexander; Hilgers, Fabienne; Loeschcke, Anita; Jaeger, Karl-Erich et al. (2016): Comparative Single-Cell Analysis of Different E. coli Expression Systems during Microfluidic Cultivation. In PloS one 11 (8), e0160711. DOI: 10.1371/journal.pone.0160711.

Bjork, Sara M.; Sjostrom, Staffan L.; Andersson-Svahn, Helene; Joensson, Haakan N. (2015): Metabolite profiling of microfluidic cell culture conditions for droplet based screening. In Biomicrofluidics 9 (4), p. 44128. DOI: 10.1063/1.4929520.

Burmeister, Alina; Hilgers, Fabienne; Langner, Annika; Westerwalbesloh, Christoph; Kerkhoff, Yannic; Tenhaef, Niklas et al. (2018): A microfluidic co-cultivation platform to investigate microbial interactions at defined microenvironments. In Lab on a chip 19 (1), pp. 98–110. DOI: 10.1039/c8lc00977e.

Cadart, Clotilde; Monnier, Sylvain; Grilli, Jacopo; Sáez, Pablo J.; Srivastava, Nishit; Attia, Rafaele et al. (2018): Size control in mammalian cells involves modulation of both growth rate and cell cycle duration. In Nature communications 9 (1), p. 3275. DOI: 10.1038/s41467-018-05393-0.

Comesaña, José F.; Otero, Juan J.; García, Emilio; Correa, Antonio (2003): Densities and Viscosities of Ternary Systems of Water + Glucose + Sodium Chloride at Several Temperatures. In J. Chem. Eng. Data 48 (2), pp. 362–366. DOI: 10.1021/je020153x.

Delvigne, Frank; Goffin, Philippe (2014): Microbial heterogeneity affects bioprocess robustness: Dynamic single-cell analysis contributes to understanding of microbial populations. In Biotechnology journal 9 (1), pp. 61–72. DOI: 10.1002/biot.201300119.

Dettinger, Philip; Frank, Tino; Etzrodt, Martin; Ahmed, Nouraiz; Reimann, Andreas; Trenzinger, Christoph et al. (2018): Automated Microfluidic System for Dynamic Stimulation and Tracking of Single Cells. In Analytical chemistry 90 (18), pp. 10695–10700. DOI: 10.1021/acs.analchem.8b00312.

Di Carlo, Dino; Wu, Liz Y.; Lee, Luke P. (2006): Dynamic single cell culture array. In Lab on a chip 6 (11), pp. 1445–1449. DOI: 10.1039/b605937f.

Du, Guansheng; Fang, Qun; den Toonder, Jaap M. J. (2016): Microfluidics for cell-based high throughput screening platforms - A review. In Analytica chimica acta 903, pp. 36–50. DOI: 10.1016/j.aca.2015.11.023.

Fritzsch, Frederik S. O.; Rosenthal, Katrin; Kampert, Anna; Howitz, Steffen; Dusny, Christian; Blank, Lars M.; Schmid, Andreas (2013): Picoliter nDEP traps enable time-resolved contactless single bacterial cell analysis in controlled microenvironments. In Lab on a chip 13 (3), pp. 397–408. DOI: 10.1039/c2lc41092c.

Gao, Yan; Li, Peng; Pappas, Dimitri (2013): A microfluidic localized, multiple cell culture array using vacuum actuated cell seeding: integrated anticancer drug testing. In Biomedical microdevices 15 (6), pp. 907–915. DOI: 10.1007/s10544-013-9779-3.

Grilo, Antonio L.; Mantalaris, Athanasios (2019): Apoptosis: A mammalian cell bioprocessing perspective. In Biotechnology advances 37 (3), pp. 459–475. DOI: 10.1016/j.biotechadv.2019.02.012.

Grünberger, Alexander; Probst, Christopher; Helfrich, Stefan; Nanda, Arun; Stute, Birgit; Wiechert, Wolfgang et al. (2015): Spatiotemporal microbial single-cell analysis using a high-throughput microfluidics cultivation platform. In Cytometry Part A 87 (12), pp. 1101–1115. DOI: 10.1002/cyto.a.22779.

Grünberger, Alexander; Probst, Christopher; Heyer, Antonia; Wiechert, Wolfgang; Frunzke, Julia; Kohlheyer, Dietrich (2013a): Microfluidic picoliter bioreactor for microbial single-cell analysis: fabrication, system setup, and operation. In Journal of visualized experiments: JoVE (82), p. 50560. DOI: 10.3791/50560.

Grünberger, Alexander; van Ooyen, Jan; Paczia, Nicole; Rohe, Peter; Schiendzielorz, Georg; Eggeling, Lothar et al. (2013b): Beyond growth rate 0.6: Corynebacterium glutamicum cultivated in highly diluted environments. In Biotechnology and bioengineering 110 (1), pp. 220–228. DOI: 10.1002/bit.24616.

Grünberger, Alexander; Wiechert, Wolfgang; Kohlheyer, Dietrich (2014): Single-cell microfluidics: opportunity for bioprocess development. In Current opinion in biotechnology 29, pp. 15–23. DOI: 10.1016/j.copbio.2014.02.008.

Halldorsson, Skarphedinn; Lucumi, Edinson; Gómez-Sjöberg, Rafael; Fleming, Ronan M. T. (2015): Advantages and challenges of microfluidic cell culture in polydimethylsiloxane devices. In Biosensors & bioelectronics 63, pp. 218–231. DOI: 10.1016/j.bios.2014.07.029.

Karakas, Hacer Ezgi; Kim, Junyoung; Park, Juhee; Oh, Jung Min; Choi, Yongjun; Gozuacik, Devrim; Cho, Yoon-Kyoung (2017): A microfluidic chip for screening individual cancer cells via eavesdropping on autophagy-inducing crosstalk in the stroma niche. In Scientific reports 7 (1), p. 2050. DOI: 10.1038/s41598-017-02172-7.

Kolnik, Martin; Tsimring, Lev S.; Hasty, Jeff (2012): Vacuum-assisted cell loading enables shear-free mammalian microfluidic culture. In Lab on a chip 12 (22), pp. 4732–4737. DOI: 10.1039/c2lc40569e.

Le, Huong; Kabbur, Santosh; Pollastrini, Luciano; Sun, Ziran; Mills, Keri; Johnson, Kevin et al. (2012): Multivariate analysis of cell culture bioprocess data--lactate consumption as process indicator. In Journal of Biotechnology 162 (2-3), pp. 210–223. DOI: 10.1016/j.jbiotec.2012.08.021.

Lecault, Véronique; White, Adam K.; Singhal, Anupam; Hansen, Carl L. (2012): Microfluidic single cell analysis: from promise to practice. In Current opinion in chemical biology 16 (3-4), pp. 381–390. DOI: 10.1016/j.cbpa.2012.03.022.

Li, Rui; Lv, Xuefei; Zhang, Xingjian; Saeed, Omer; Deng, Yulin (2016): Microfluidics for cell-cell interactions: A review. In Front. Chem. Sci. Eng. 10 (1), pp. 90–98. DOI: 10.1007/s11705-015-1550-2.

Lin, Ching-Hui; Hsiao, Yi-Hsing; Chang, Hao-Chen; Yeh, Chuan-Feng; He, Cheng-Kun; Salm, Eric M. et al. (2015): A microfluidic dual-well device for high-throughput single-cell capture and culture. In Lab on a chip 15 (14), pp. 2928–2938. DOI: 10.1039/c5lc00541h.

Lindström, Sara; Andersson-Svahn, Helene (2010): Overview of single-cell analyses: microdevices and applications. In Lab on a chip 10 (24), pp. 3363–3372. DOI: 10.1039/c0lc00150c.

Luni, Camilla; Giulitti, Stefano; Serena, Elena; Ferrari, Luca; Zambon, Alessandro; Gagliano, Onelia et al. (2016): High-efficiency cellular reprogramming with microfluidics. In Nature methods 13 (5), pp. 446–452. DOI: 10.1038/nmeth.3832.

Marques, Marco Pc; Szita, Nicolas (2017): Bioprocess microfluidics: applying microfluidic devices for bioprocessing. In Current opinion in chemical engineering 18, pp. 61–68. DOI: 10.1016/j.coche.2017.09.004.

Mehling, Matthias; Tay, Savaş (2014): Microfluidic cell culture. In Current opinion in biotechnology 25, pp. 95–102. DOI: 10.1016/j.copbio.2013.10.005.

Nolan, Ryan P.; Lee, Kyongbum (2011): Dynamic model of CHO cell metabolism. In Metabolic engineering 13 (1), pp. 108–124. DOI: 10.1016/j.ymben.2010.09.003.

Phoenix, David (1997): Introductory mathematics for the life sciences. Boca Raton, Fla.: CRC Press (Modules in life sciences).

Potapova, Tamara A.; Daum, John R.; Pittman, Bradley D.; Hudson, Joanna R.; Jones, Tara N.; Satinover, David L. et al. (2006): The reversibility of mitotic exit in vertebrate cells. In Nature 440 (7086), pp. 954–958. DOI: 10.1038/nature04652.

Raimes, William; Rubi, Mathieu; Super, Alexandre; Marques, Marco P. C.; Veraitch, Farlan; Szita, Nicolas (2017): Transfection in perfused microfluidic cell culture devices: A case study. In Process biochemistry 59 (Pt B), pp. 297–302. DOI: 10.1016/j.procbio.2016.09.006.

Rowat, Amy C.; Bird, James C.; Agresti, Jeremy J.; Rando, Oliver J.; Weitz, David A. (2009): Tracking lineages of single cells in lines using a microfluidic device. In Proceedings of the National Academy of Sciences of the United States of America 106 (43), pp. 18149–18154. DOI: 10.1073/pnas.0903163106.

Schindelin, Johannes; Arganda-Carreras, Ignacio; Frise, Erwin; Kaynig, Verena; Longair, Mark; Pietzsch, Tobias et al. (2012): Fiji: an open-source platform for biological-image analysis. In Nature methods 9 (7), pp. 676–682. DOI: 10.1038/nmeth.2019.

Schmitz, Julian; Noll, Thomas; Grünberger, Alexander (2019): Heterogeneity Studies of Mammalian Cells for Bioproduction: From Tools to Application. In Trends in biotechnology 37 (6), pp. 645–660. DOI: 10.1016/j.tibtech.2018.11.007.

Skylar-Scott, Mark A.; Uzel, Sebastien G. M.; Nam, Lucy L.; Ahrens, John H.; Truby, Ryan L.; Damaraju, Sarita; Lewis, Jennifer A. (2019): Science Journals — AAAS // Biomanufacturing of organ-specific tissues with high cellular density and embedded vascular channels. In Science advances 5 (9), eaaw2459. DOI: 10.1126/sciadv.aaw2459.

Taheri-Araghi, Sattar; Brown, Steven D.; Sauls, John T.; McIntosh, Dustin B.; Jun, Suckjoon (2015): Single-Cell Physiology. In Annual review of biophysics 44, pp. 123–142. DOI: 10.1146/annurev-biophys-060414-034236.

Täuber, Sarah; Lieres, Eric von; Grünberger, Alexander (2020): Dynamic Environmental Control in Microfluidic Single-Cell Cultivations: From Concepts to Applications. In Small (Weinheim an der Bergstrasse, Germany), e1906670. DOI: 10.1002/smll.201906670.

Templer, Richard H.; Ces, Oscar (2008): New frontiers in single-cell analysis. In Journal of the Royal Society, Interface 5 Suppl 2, S111–2. DOI: 10.1098/rsif.2008.0279.focus.

Unthan, Simon; Grünberger, Alexander; van Ooyen, Jan; Gätgens, Jochem; Heinrich, Johanna; Paczia, Nicole et al. (2014): Beyond growth rate 0.6: What drives Corynebacterium glutamicum to higher growth rates in defined medium. In Biotechnology and bioengineering 111 (2), pp. 359–371. DOI: 10.1002/bit.25103.

Vuaridel-Thurre, Gaëlle; Vuaridel, Ambroise R.; Dhar, Neeraj; McKinney, John D. (2020): Computational Analysis of the Mutual Constraints between Single-Cell Growth and Division Control Models. In Advanced biosystems 4 (2), e1900103. DOI: 10.1002/adbi.201900103.

Walsh, Gary (2018): Biopharmaceutical benchmarks 2018. In Nature biotechnology 36 (12), pp. 1136–1145. DOI: 10.1038/nbt.4305.

Wang, Daojing; Bodovitz, Steven (2010): Single cell analysis: the new frontier in ‘omics’. In Trends in biotechnology 28 (6), pp. 281–290. DOI: 10.1016/j.tibtech.2010.03.002.

Westerwalbesloh, Christoph; Grünberger, Alexander; Stute, Birgit; Weber, Sophie; Wiechert, Wolfgang; Kohlheyer, Dietrich; Lieres, Eric von (2015): Modeling and CFD simulation of nutrient distribution in picoliter bioreactors for bacterial growth studies on single-cell level. In Lab on a chip 15 (21), pp. 4177–4186. DOI: 10.1039/c5lc00646e.

Westerwalbesloh, Christoph; Grünberger, Alexander; Wiechert, Wolfgang; Kohlheyer, Dietrich; Lieres, Eric von (2017): Coarse-graining bacteria colonies for modelling critical solute distributions in picolitre bioreactors for bacterial studies on single-cell level. In Microbial biotechnology 10 (4), pp. 845–857. DOI: 10.1111/1751-7915.12708.

Wheeler, Aaron R.; Throndset, William R.; Whelan, Rebecca J.; Leach, Andrew M.; Zare, Richard N.; Liao, Yish Hann et al. (2003): Microfluidic device for single-cell analysis. In Analytical chemistry 75 (14), pp. 3581–3586. DOI: 10.1021/ac0340758.

Wurm, Florian M. (2004): Production of recombinant protein therapeutics in cultivated mammalian cells. In Nature biotechnology 22 (11), pp. 1393–1398. DOI: 10.1038/nbt1026.

Yin, Huabing; Marshall, Damian (2012): Microfluidics for single cell analysis. In Current opinion in biotechnology 23 (1), pp. 110–119. DOI: 10.1016/j.copbio.2011.11.002.

