## Supplementary Information for "Development and application of a cultivation platform for mammalian suspension cell lines with single-cell resolution (MaSC)"

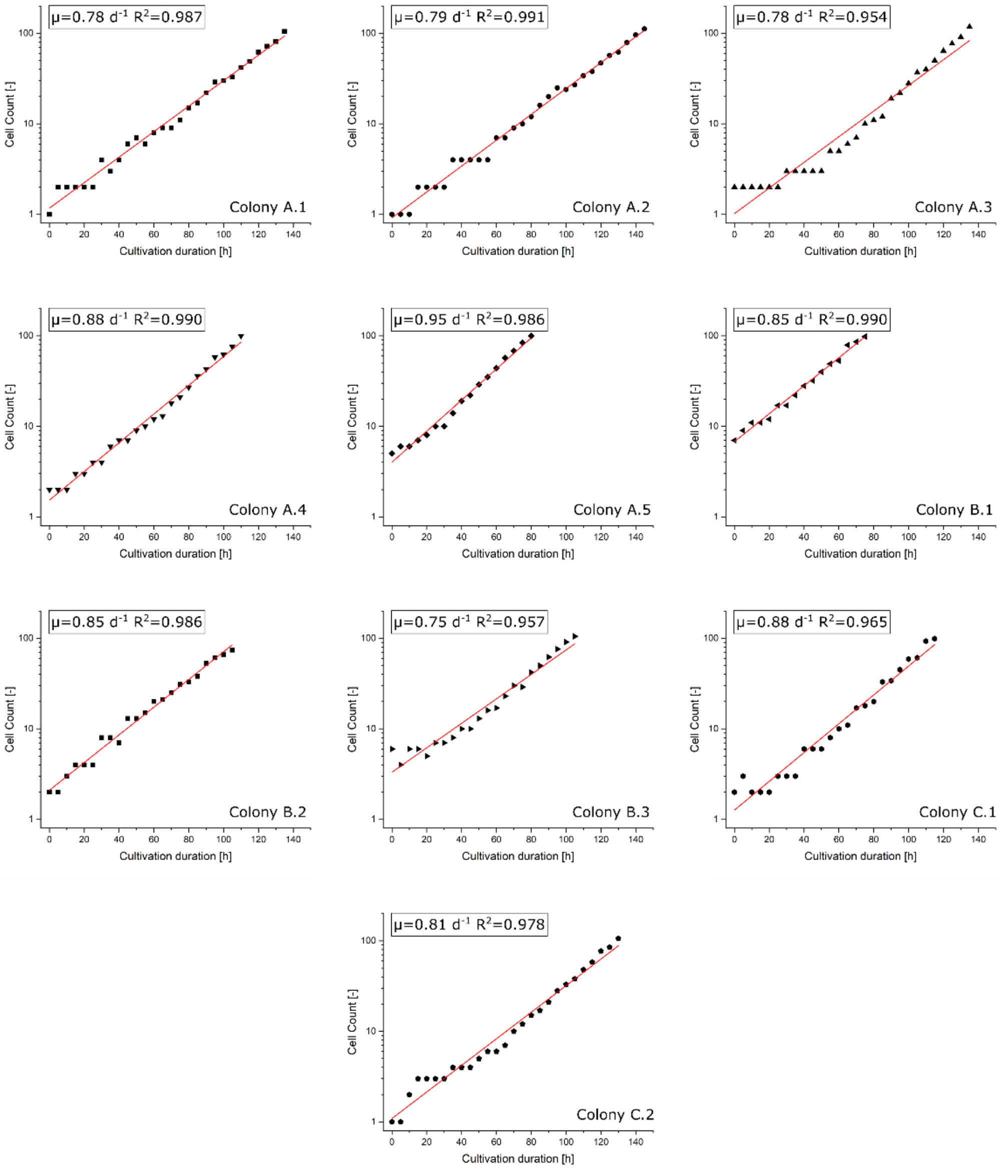


Fig. S1: Semi-logarithmically plotted growth profile of the analyzed colonies with their resulting linear fit. For every colony the graphically determined growth rate μ with its coefficient of determination $R^2$ are placed in the upper part of the respective plot.

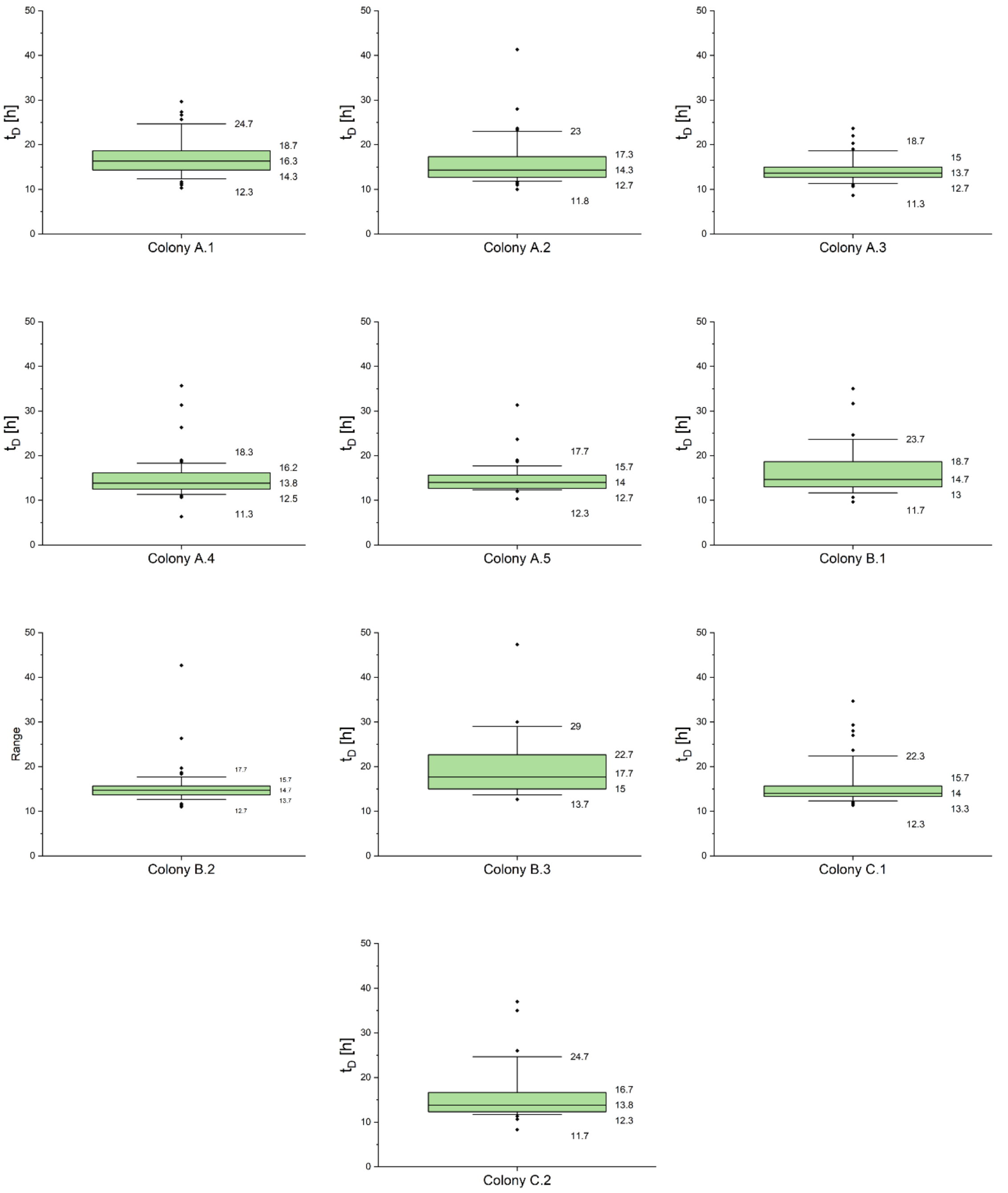


Fig. S2: Distribution of single-cell doubling times of the ten studied colonies. The colored segment marks the interquartile range from 25% to 75 % and the horizontal line the median. The whiskers represent the 10 % and 90 % percentile and the tilted squares mark rare cellular events. Values for interquartile boards and whiskers are depicted.

Tab. S1: Comparison of the growth rates and doubling times depicted from colony level and single-cell level. $\mu_{colony}$ was transformed into $t_{D,\ colony}$ and $t_{D,\ single\text{-}cell}$ was transformed into $\mu_{single\text{-}cell}$ using equation (1).

| | $\mu_{colony}$ | $t_{D,\ colony}$ | $\mu_{single\text{-}cell}$ | $t_{D,\ single\text{-}cell}$ |
|---|---|---|---|---|
| Colony A.1 | 0.78 | 21.33 | 1.00 | 16.71 |
| Colony A.2 | 0.79 | 21.06 | 1.08 | 15.40 |
| Colony A.3 | 0.78 | 21.33 | 1.19 | 14.02 |
| Colony A.4 | 0.88 | 18.90 | 1.15 | 14.41 |
| Colony A.5 | 0.95 | 17.51 | 1.14 | 14.57 |
| Colony B.1 | 0.85 | 19.57 | 1.05 | 15.77 |
| Colony B.2 | 0.85 | 19.57 | 1.10 | 15.10 |
| Colony B.3 | 0.75 | 22.18 | 1.00 | 18.68 |
| Colony C.1 | 0.88 | 18.90 | 1.10 | 15.09 |
| Colony C.2 | 0.81 | 20.54 | 1.10 | 15.11 |

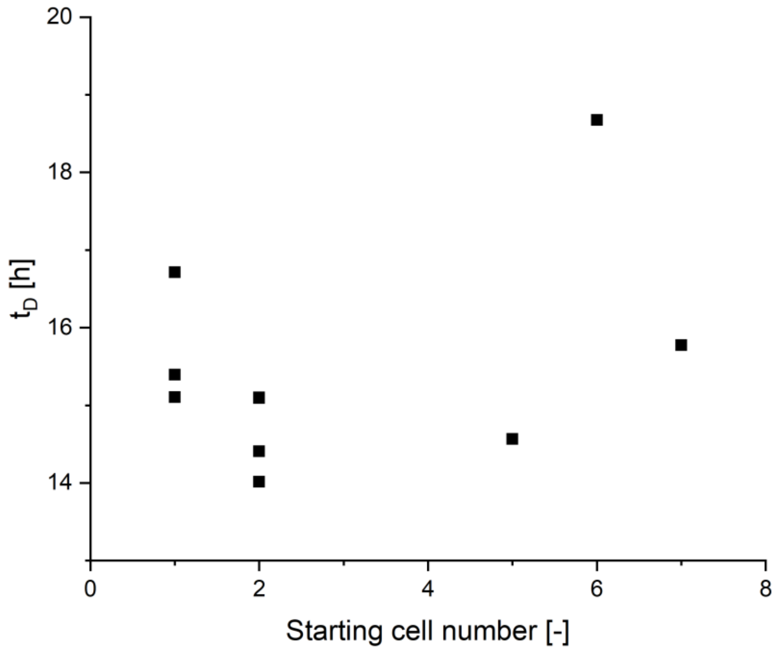


Fig. S3: Correlation between starting cell count of each of all ten colonies (Fig. S1) and the mean doubling time $t_D$ (Fig. S2). A correlation coefficient of R = 0.394 shows only weak positive correlation between initial cell number and mean doubling time, therefore initial cell number only influence the mean doubling time of the respective colony weakly. CHO K1 suspension cells were cultivated under MaSC conditions as described in ‘Materials and Methods’.

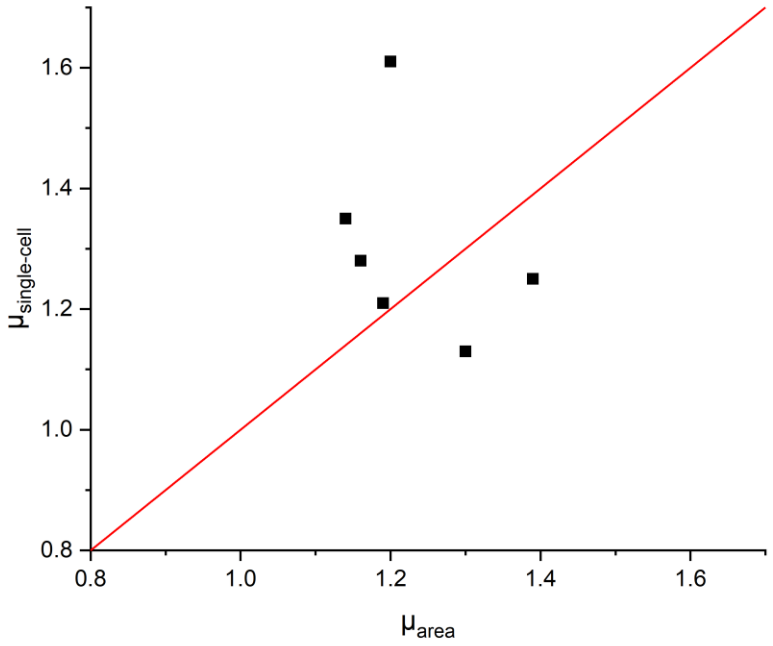


Fig. S4: Correlation between the area related single-cell growth rate $\mu_{area}$ (Fig. 5) and the growth rate $\mu_{single\text{-}cell}$ resulting from the individual doubling times $t_D$ of the selected single cells from colony A.5. CHO K1 suspension cells were cultivated under MaSC conditions as described in 'Materials and Methods'.

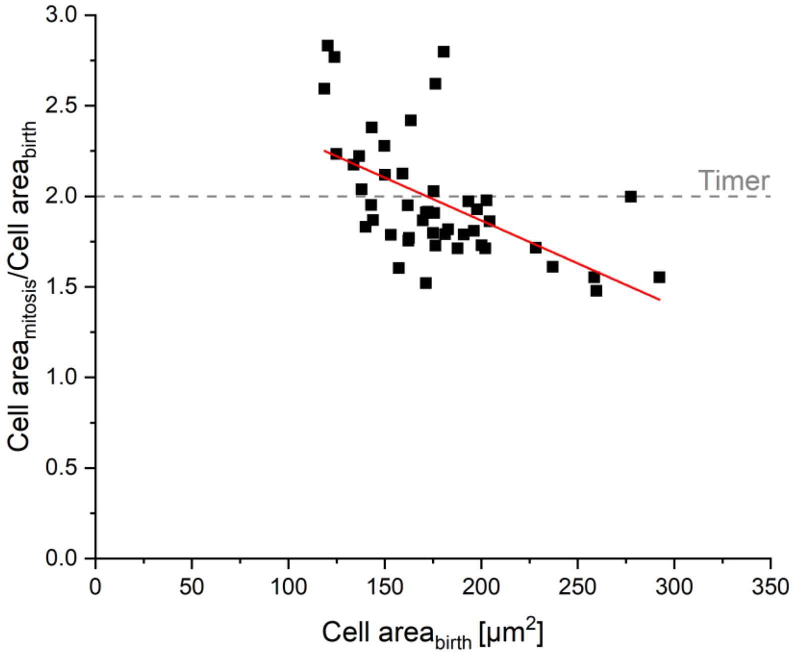


Fig. S5: Cell size homeostasis analysis of single cells from colony A.1 (Fig. 4 B). To investigate the agreement of CHO K1 suspension cells with the timer cell size homeostasis model, the correlation between cell area at birth and the ratio of cell area at mitosis and cell area at birth was analyzed. The dashed grey line shows the expected trend in case of a timer behavior.
